# Plastic-degrading clusters of orthologous groups reveal near-universal biodegradation potential in prokaryotes

**DOI:** 10.1101/2025.06.10.658834

**Authors:** Shakira Mustari, Loan Tú Phạm, Kari Saikkonen, Miho Nakamura, Pere Puigbò

## Abstract

Micro- and nanoplastic pollution (MNPP) is an increasing environmental threat due to the persistence, dispersal, and potential toxicity of plastic particles. Although microbial biodegradation offers a sustainable mitigation strategy, a comprehensive understanding of plastic-degrading proteins across microbial taxa is lacking. Here, we present the Plastic-Degrading Clusters of Orthologous Groups (PDCOGs) database (https://phylobone.com/microworld/PDCOG), comprising 625,616 potential plastic-degrading proteins (PPDPs) from free-living prokaryotes organized into 51 orthologous groups. The database/PDCOGs enable systematic analysis of microbial plastic-degrading capacity across ecosystems and phylogenetic lineages. Notably, PPDPs constitute ∼3.5% of all prokaryotic proteins, with over 95% of the species having the potential to biodegrade at least one plastic polymer type. This resource provides a genomic tool/framework for exploring the ecological and evolutionary importance of plastic biodegradation and supports future efforts to mitigate the global MNPP crisis.

## MAIN

Plastic pollution is among the most urgent environmental challenges of the 21st century owing to its small size, persistence in the environment, dispersal, and potential adverse effects on ecosystems and human health ^1^. Microplastics (1–1000 µm) and nanoplastics (0.001–1 µm) ^2^ arise from the fragmentation of larger plastic items or are directly manufactured at small scales ^3^. They have been detected across diverse habitats, including marine, freshwater, terrestrial and atmospheric environments, where they pose serious threats to wildlife and may enter the human food chain ^4,5^. In addition to direct toxicity, micro- and nanoplastics (MNPs) can serve as vectors for chemical pollutants and pathogenic microbes, raising concerns about the indirect health impacts on humans when they consume contaminated food and water ^6,7^. Recent studies have also shown that MNPs may play an important role in the dissemination of antimicrobial-resistant strains because of their capacity to host complex microbial communities and antimicrobial resistance genes through biofilm formation ^8^. In addition, nanoplastics seem to have a more negative impact on ecosystems ^9^ and health ^10^, than microplastics do because of their greater reactivity, abundance, and ability to cross barriers and accumulate in tissues.

Climate change is expected to accelerate MNP accumulation across the Earth. Globally, it is estimated that over 14 million tons of microplastics have accumulated on the seafloor ^11^. Despite increasing awareness, current strategies to mitigate this MNP pollution problem remain fragmented and largely ineffective in offering long-term solutions. The ubiquity of MNPs, which affect both the most densely populated regions and remote environments such as the Arctic and Antarctic, underscores the global scale of this issue ^12,13^. Moreover, the interaction between plastic pollution and climate warming presents an emerging threat ^14^. Ocean currents transport plastic debris to polar regions, while melting sea ice releases previously trapped microplastics ^12,13^. The Arctic Ocean is predicted to be ice-free before the end of the century ^15^, whereas ice-free areas in Antarctica may expand from the present 2% to nearly 25% in the same time frame ^16^. These changes will likely exacerbate plastic accumulation in these ecologically vulnerable regions.

Despite increasing awareness, current strategies for managing plastic waste remain fragmented and ineffective in offering long-term solutions at the global scale. Given the persistence of common plastic polymers, such as polyethylene (PE), polystyrene (PS), polypropylene (PP), polyvinyl chloride (PVC), polyethylene terephthalate (PET), and polyurethane (PUR), microbial biodegradation has emerged as a promising, sustainable and environmentally friendly alternative ^17^. Studies of the plastisphere—the microbial biofilms that are formed on plastic surfaces ^18^—have revealed a growing diversity of microbial strains capable of plastic degradation ^19^. Although earlier studies focused primarily on mesophilic bacteria (20–45 °C) ^20^, a recent study identified plastic-degrading enzymes from thermophiles (i.e., microbial strains with optimal growth temperatures above 45 °C ^20^). Thermostable enzymes are likely more suitable for industrial applications ^21^. Furthermore, microbial strains isolated from the Arctic plastisphere have demonstrated the ability to break down biodegradable plastic films at temperatures below 20 °C ^22^, which may offer increased stability and efficiency for industrial and cold-environment applications ^21^. However, despite these advances, the global understanding of plastic-degrading microbial capacities remains limited by the absence of systematic, large-scale comparative resources.

To address this gap, we introduce the Plastic-Degrading Clusters of Orthologous Groups (PDCOGs) database—a comprehensive resource of putative plastic-degrading proteins (PPDPs) from free-living bacteria allocated into 51 orthologous groups. The PDCOGs database enables systematic comparisons of PPDPs across environmental gradients and geographic regions, facilitating a deeper understanding of the global biodegradation landscape. Ultimately, this resource provides a foundation for the development of microbial and enzymatic solutions to mitigate the growing threat of MNP pollution under rapidly changing environmental conditions.

## RESULTS

### A global resource of putative plastic-degrading protein clusters

We present PDCOGs, a database of plastic-degrading clusters of orthologous groups in prokaryotes that is freely available at http://phylobone.com/microworld/PDCOG. The resource is organized into six sections and comprises 662,830 PPDP sequences categorized into 51 orthologous clusters (PDCOGs) (Table 1, Supplementary Table S1). The PPDPs span diverse environmental sources and are associated with enzymatic activities involved in the biodegradation of both natural and synthetic polymers.

**Table 1.** List of PDCOGS and their distributions across prokaryotic species, plastic types and environments.

| PDCOG | Number | COG | Fun | %Prok | Plastic types | Environmental distribution |
| --- | --- | --- | --- | --- | --- | --- |
| PDCOG001 | 20,071 | COG0154 | J | 88.81% | N: 0/11; S_He: 1/21; S_Ho: 0/7 | Ai (0.56%), Aq (60.27%), Te (38.97%), Un (0.2%) |
| PDCOG002 | 21,153 | COG0277 | C | 80.49% | N: 0/11; S_He: 2/21; S_Ho: 1/7 | Ai (0.54%), Aq (56.41%), Te (42.84%), Un (0.21%) |
| PDCOG003 | 6,475 | COG0400 | R | 52.44% | N: 0/11; S_He: 1/21; S_Ho: 0/7 | Ai (0.59%), Aq (73.27%), Te (26.01%), Un (0.14%) |
| PDCOG004 | 12,730 | COG0412 | Q | 57.01% | N: 2/11; S_He: 9/21; S_Ho: 1/7 | Ai (0.46%), Aq (53.94%), Te (45.48%), Un (0.13%) |
| PDCOG005 | 16,725 | COG0508 | C | 80.01% | N: 0/11; S_He: 1/21; S_Ho: 1/7 | Ai (0.50%), Aq (68.55%), Te (30.75%), Un (0.2%) |
| PDCOG006 | 90,850 | COG0596 | HR | 91.29% | N: 4/11; S_He: 8/21; S_Ho: 1/7 | Ai (0.56%), Aq (54.26%), Te (45.01%), Un (0.16%) |
| PDCOG007 | 4,527 | COG0627 | V | 43.07% | N: 1/11; S_He: 5/21; S_Ho: 1/7 | Ai (0.46%), Aq (48.13%), Te (51.25%), Un (0.15%) |
| PDCOG008 | 15,526 | COG0657 | I | 65.11% | N: 3/11; S_He: 6/21; S_Ho: 2/7 | Ai (0.71%), Aq (51.13%), Te (48.04%), Un (0.12%) |
| PDCOG009 | 14,723 | COG0726 | MG | 80.66% | N: 0/11; S_He: 0/21; S_Ho: 2/7 | Ai (0.60%), Aq (51.69%), Te (47.48%), Un (0.22%) |
| PDCOG010 | 43,441 | COG0834 | TE | 76.96% | N: 0/11; S_He: 1/21; S_Ho: 0/7 | Ai (0.46%), Aq (54.29%), Te (45.04%), Un (0.22%) |
| PDCOG011 | 67,608 | COG1012 | I | 89.46% | N: 0/11; S_He: 1/21; S_Ho: 1/7 | Ai (0.44%), Aq (53.08%), Te (46.25%), Un (0.22%) |
| PDCOG012 | 139,666 | COG1028 | I | 95.03% | N: 3/11; S_He: 0/21; S_Ho: 2/7 | Ai (0.57%), Aq (55.03%), Te (44.21%), Un (0.19%) |
| PDCOG013 | 4,116 | COG1073 | T | 30.31% | N: 1/11; S_He: 5/21; S_Ho: 1/7 | Ai (0.56%), Aq (65.38%), Te (33.79%), Un (0.27%) |
| PDCOG014 | 2,434 | COG1075 | I | 32.58% | N: 4/11; S_He: 8/21; S_Ho: 1/7 | Ai (0.58%), Aq (49.92%), Te (49.30%), Un (0.21%) |
| PDCOG015 | 261 | COG1382 | O | 8.89% | N: 0/11; S_He: 1/21; S_Ho: 0/7 | Ai (1.53%), Aq (69.35%), Te (29.12%), Un (0.00%) |
| PDCOG016 | 4,483 | COG1398 | I | 27.57% | N: 0/11; S_He: 0/21; S_Ho: 1/7 | Ai (0.27%), Aq (61.25%), Te (38.39%), Un (0.09%) |
| PDCOG017 | 6,857 | COG1404 | O | 62.37% | N: 1/11; S_He: 4/21; S_Ho: 0/7 | Ai (0.61%), Aq (54.53%), Te (44.55%), Un (0.31%) |
| PDCOG018 | 9,776 | COG1506 | E | 71.65% | N: 4/11; S_He: 4/21; S_Ho: 0/7 | Ai (0.30%), Aq (71.69%), Te (27.91%), Un (0.11%) |
| PDCOG019 | 2,783 | COG1647 | Q | 31.84% | N: 1/11; S_He: 5/21; S_Ho: 1/7 | Ai (0.25%), Aq (63.71%), Te (35.79%), Un (0.25%) |
| PDCOG020 | 21,699 | COG1680 | V | 66.77% | N: 1/11; S_He: 5/21; S_Ho: 0/7 | Ai (0.41%), Aq (59.08%), Te (40.36%), Un (0.15%) |
| PDCOG021 | 4,746 | COG1773 | P | 42.55% | N: 0/11; S_He: 0/21; S_Ho: 3/7 | Ai (0.36%), Aq (62.45%), Te (37.04%), Un (0.15%) |
| PDCOG022 | 4,733 | COG1858 | O | 35.76% | N: 0/11; S_He: 0/21; S_Ho: 2/7 | Ai (0.42%), Aq (58.27%), Te (41.12%), Un (0.19%) |
| PDCOG023 | 18,460 | COG2010 | C | 56.58% | N: 0/11; S_He: 0/21; S_Ho: 2/7 | Ai (0.64%), Aq (56.21%), Te (43.01%), Un (0.14%) |
| PDCOG024 | 6,773 | COG2132 | DMP | 51.09% | N: 0/11; S_He: 0/21; S_Ho: 4/7 | Ai (0.75%), Aq (56.18%), Te (42.85%), Un (0.22%) |
| PDCOG025 | 590 | COG2247 | M | 3.44% | N: 0/11; S_He: 1/21; S_Ho: 0/7 | Ai (0.34%), Aq (68.98%), Te (30.68%), Un (0.00%) |
| PDCOG026 | 8,632 | COG2267 | I | 70.86% | N: 3/11; S_He: 14/21; S_Ho: 0/7 | Ai (0.47%), Aq (57.75%), Te (41.52%), Un (0.25%) |
| PDCOG027 | 2,972 | COG2272 | I | 20.17% | N: 1/11; S_He: 9/21; S_Ho: 0/7 | Ai (0.34%), Aq (63.36%), Te (36.07%), Un (0.24%) |
| PDCOG028 | 17,646 | COG2303 | IR | 49.96% | N: 0/11; S_He: 1/21; S_Ho: 1/7 | Ai (0.49%), Aq (56.58%), Te (42.76%), Un (0.17%) |
| PDCOG029 | 1,542 | COG2703 | T | 20.82% | N: 0/11; S_He: 0/21; S_Ho: 1/7 | Ai (0.71%), Aq (66.67%), Te (32.10%), Un (0.52%) |
| PDCOG030 | 10,074 | COG2863 | C | 35.37% | N: 0/11; S_He: 1/21; S_Ho: 1/7 | Ai (0.60%), Aq (59.90%), Te (39.40%), Un (0.11%) |
| PDCOG031 | 5,066 | COG2931 | Q | 28.75% | N: 0/11; S_He: 2/21; S_Ho: 0/7 | Ai (0.71%), Aq (61.94%), Te (36.89%), Un (0.45%) |
| PDCOG032 | 3,397 | COG3145 | L | 33.97% | N: 0/11; S_He: 0/21; S_Ho: 3/7 | Ai (0.71%), Aq (59.58%), Te (39.48%), Un (0.24%) |
| PDCOG033 | 2,814 | COG3147 | D | 28.22% | N: 2/11; S_He: 0/21; S_Ho: 0/7 | Ai (0.39%), Aq (66.24%), Te (33.08%), Un (0.28%) |
| PDCOG034 | 1,515 | COG3179 | G | 11.80% | N: 0/11; S_He: 1/21; S_Ho: 0/7 | Ai (0.20%), Aq (43.17%), Te (56.44%), Un (0.20%) |
| PDCOG035 | 2,522 | COG3191 | E | 29.05% | N: 0/11; S_He: 2/21; S_Ho: 0/7 | Ai (0.36%), Aq (42.27%), Te (57.14%), Un (0.24%) |
| PDCOG036 | 1,0447 | COG3239 | I | 40.29% | N: 0/11; S_He: 0/21; S_Ho: 4/7 | Ai (0.30%), Aq (61.02%), Te (38.57%), Un (0.11%) |
| PDCOG037 | 3,984 | COG3243 | I | 28.83% | N: 2/11; S_He: 0/21; S_Ho: 0/7 | Ai (0.85%), Aq (56.58%), Te (42.42%), Un (0.15%) |
| PDCOG038 | 693 | COG3266 | D | 17.77% | N: 2/11; S_He: 0/21; S_Ho: 0/7 | Ai (0.43%), Aq (82.40%), Te (16.74%), Un (0.43%) |
| PDCOG039 | 1,892 | COG3325 | G | 16.33% | N: 0/11; S_He: 4/21; S_Ho: 0/7 | Ai (0.16%), Aq (53.91%), Te (45.61%), Un (0.32%) |
| PDCOG040 | 278 | COG3401 | R | 11.28% | N: 0/11; S_He: 4/21; S_Ho: 0/7 | Ai (0.36%), Aq (57.91%), Te (41.73%), Un (0.00%) |
| PDCOG041 | 3,327 | COG3509 | G | 25.04% | N: 8/11; S_He: 11/21; S_Ho: 2/7 | Ai (0.75%), Aq (53.77%), Te (45.33%), Un (0.15%) |
| PDCOG042 | 1,500 | COG3591 | O | 21.56% | N: 0/11; S_He: 1/21; S_Ho: 0/7 | Ai (0.47%), Aq (49.87%), Te (48.87%), Un (0.80%) |
| PDCOG043 | 1,294 | COG3979 | G | 11.67% | N: 0/11; S_He: 5/21; S_Ho: 0/7 | Ai (0.31%), Aq (69.78%), Te (29.75%), Un (0.15%) |
| PDCOG044 | 2,311 | COG4188 | R | 21.86% | N: 3/11; S_He: 8/21; S_Ho: 0/7 | Ai (0.17%), Aq (65.73%), Te (34.01%), Un (0.09%) |
| PDCOG045 | 2,504 | COG4553 | I | 14.94% | N: 3/11; S_He: 7/21; S_Ho: 0/7 | Ai (0.72%), Aq (38.74%), Te (60.42%), Un (0.12%) |
| PDCOG046 | 9,970 | COG1366 | T | 35.76% | N: 0/11; S_He: 1/21; S_Ho: 0/7 | Ai (0.90%), Aq (57.92%), Te (40.52%), Un (0.65%) |
| PDCOG047 | 6,948 | COG1520 | M | 53.53% | N: 0/11; S_He: 1/21; S_Ho: 0/7 | Ai (0.39%), Aq (70.31%), Te (29.07%), Un (0.23%) |
| PDCOG048 | 15,226 | COG2755 | DI | 73.78% | N: 0/11; S_He: 1/21; S_Ho: 0/7 | Ai (0.63%), Aq (57.97%), Te (41.11%), Un (0.28%) |
| PDCOG049 | 2,710 | COG2819 | R | 24.65% | N: 0/11; S_He: 1/21; S_Ho: 0/7 | Ai (0.44%), Aq (66.94%), Te (32.55%), Un (0.07%) |
| PDCOG050 | 1,749 | COG3240 | I | 12.46% | N: 0/11; S_He: 1/21; S_Ho: 0/7 | Ai (0.80%), Aq (45.45%), Te (53.57%), Un (0.17%) |
| PDCOG051 | 611 | COG4099 | R | 16.16% | N: 0/11; S_He: 1/21; S_Ho: 0/7 | Ai (0.33%), Aq (73.65%), Te (26.02%), Un (0.00%) |

The data capture the microbial degradation potential across 11 natural and 28 synthetic polymers, including 7 homochain and 21 heterochain variants. Notably, polyhydroxyalkanoates such as P(3HB-co-3MP), P(3HV) ^23^, and P3HP ^24^, although often considered natural owing to their microbial biosynthesis, and classified here as natural ^25,26^, are engineered for production in *Escherichia coli, Clostridium butiricum*, and *Klebsiella pneumoniae* ^27^. All the plastic polymers included in the PDCOGs databases are potentially susceptible to biodegradation by microbes encoding at least one copy of a PDCOG (Supplementary Datasets SD1-2).

### Linking PDCOGs with polymers and microbial taxa

A bipartite network analysis linking PDCOGs to polymer types and microbial taxa revealed substantial functional and phylogenetic diversity (Fig. 1 and Supplementary Fig. S1). This modular structure suggests that plastic-degrading potential is broadly distributed across bacterial lineages (Supplementary Dataset SD3). The median number of polymer associations per PDCOG was 3 (range: 1–16). PDCOG041 and PDCOG026 display the highest degree of connectivity and are associated with 16 and 12 polymers, respectively. In contrast, 22 PDCOGs are associated exclusively with a single polymer type, suggesting potential specificity: PDCOG003 is linked to nylon, PDCOG005 is linked to PET, PDCOG007 is linked to PET, PDCOG009 is linked to PET, PDCOG010 is linked to low-density polyethylene (LDPE), PDCOG011 is linked to PET, PDCOG015 is linked to polyethylene glycol (PEG), PDCOG016 is linked to polylactic acid (PLA), PDCOG017 is linked to LDPE, PDCOG017 is linked to PLA, PDCOG019 is linked to PET, PDCOG022 is linked to natural rubber (NR), PDCOG023 is linked to NR, PDCOG025 is linked to PLA, PDCOG028 is linked to PEG, PDCOG029 is linked to NR, PDCOG030 is linked to polyvinyl alcohol (PVA), PDCOG031 is linked to polyurethane (PU), PDCOG032 is linked to PS, PDCOG034 is linked to polycaprolactone (PCL), PDCOG035 is linked to nylon, and PDCOG042 is linked to PLA.

**Fig. 1.**
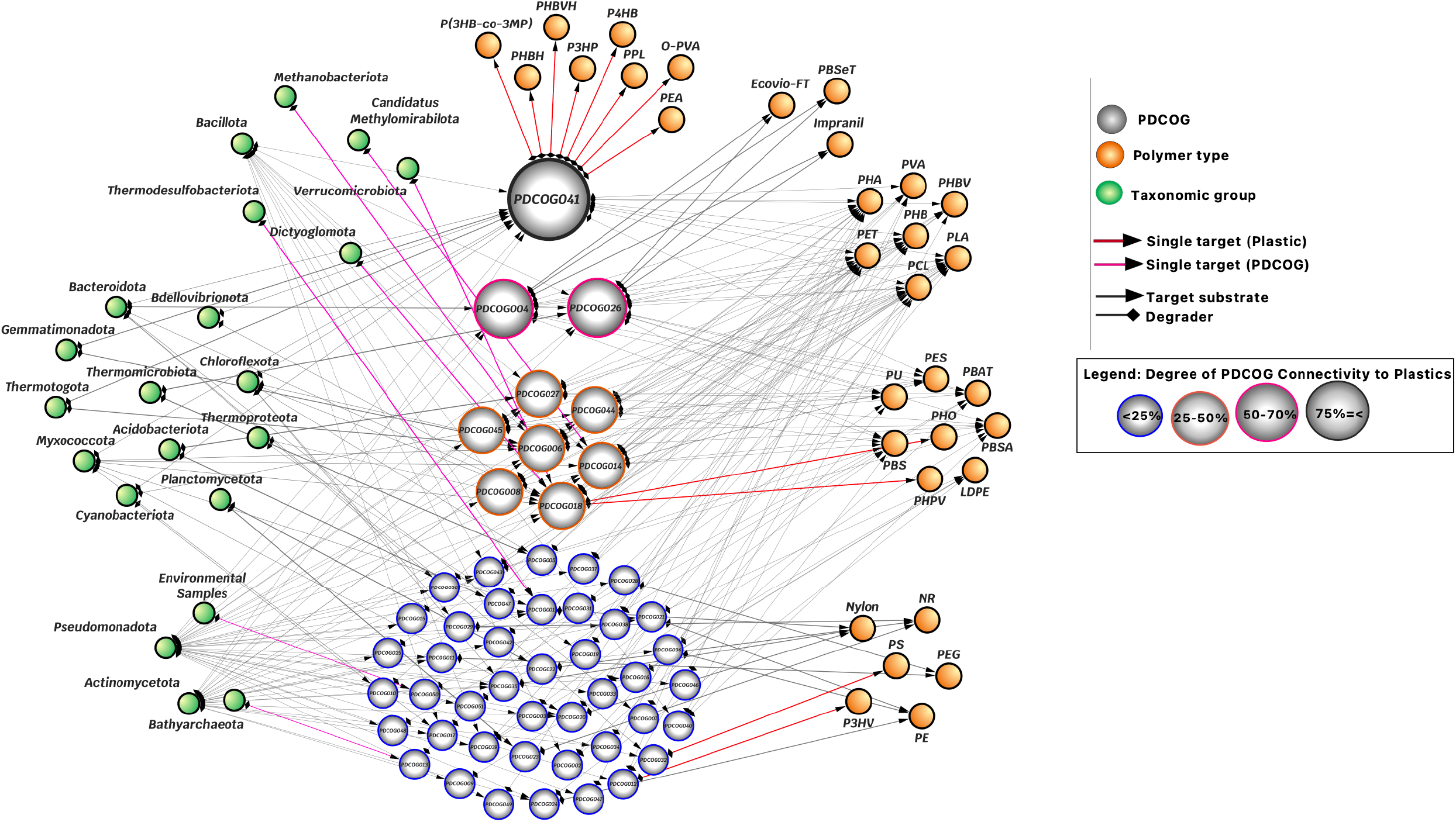
Bipartite network of associations between the plastic-degrading clusters of orthologous groups (PDCOGs) and microbial species and enzymatic functions. The network was generated from the linkage data of organisms, COGs, and targeted plastics using Cytoscape v3.10.3. In the network, node colors represent (i) the COG functional hierarchy on the basis of connectivity to polymers, (ii) the taxonomic clustering of species, and (iii) the classification of plastic polymers. Edge colors indicate the strength of connections, weighted by interaction confidence.

The distribution of microbial species in natural environments, based on the best BLAST hits of the seed dataset in the COG database, varies across PDCOGs, ranging from 1 to 56 species. PDCOG041 was associated with the most species (n = 56), followed by PDCOG006 (34), PDCOG014 (33), PDCOG018 (25), PDCOG027 (21), PDCOG045 (19), and PDCOG001 (10). The remaining PDCOGs are each associated with fewer than 10 species (Supplementary Dataset SD3). Moreover, several bacterial groups are associated with a single PDCOG, such as Thermodesulfobacteria with PDCOG001, Dictyoglomota and Verrucomicrobiota with PDCOG018, Methylomirabilota with PDCOG014, and Methanobacteriota with PDCOG006.

### General functional characterization of the PDCOGs

The 51 PDCOGs encompass a wide range of predicted biological functions. A functional characterization of the PDCOGs using the 26 functional categories (Supplementary Table S2) in the COG database revealed that the functional categories of PPDP are I (lipid transport and metabolism; n = 12), R (general function prediction only; n = 6), and G (carbohydrate transport and metabolism; n = 5), which are the most abundant among plastic-degrading enzymes, followed by the other categories in this order: C=D=M=O=T>E=Q>P=V>H=J=L (Fig. 2A). Among the 51 PDCOGs, the enzymes involved in plastic biodegradation include oxidoreductases (EC1s; 33.33%), transferases (EC2s; 33.33%), hydrolases (EC3s; 52.94%), lyases (EC4s; 5.88%), isomerases (EC5s; 1.96%), ligases (EC6s; 9.8%) and translocases (EC7s; 1.96%) (Supplementary Dataset SD4). All PDCOGs were also annotated with KEGG pathway information, gene ontology terms, and Interpro functional domains (Supplementary Dataset SD4), facilitating integrative functional and systems-level analyses.

**Fig. 2.**
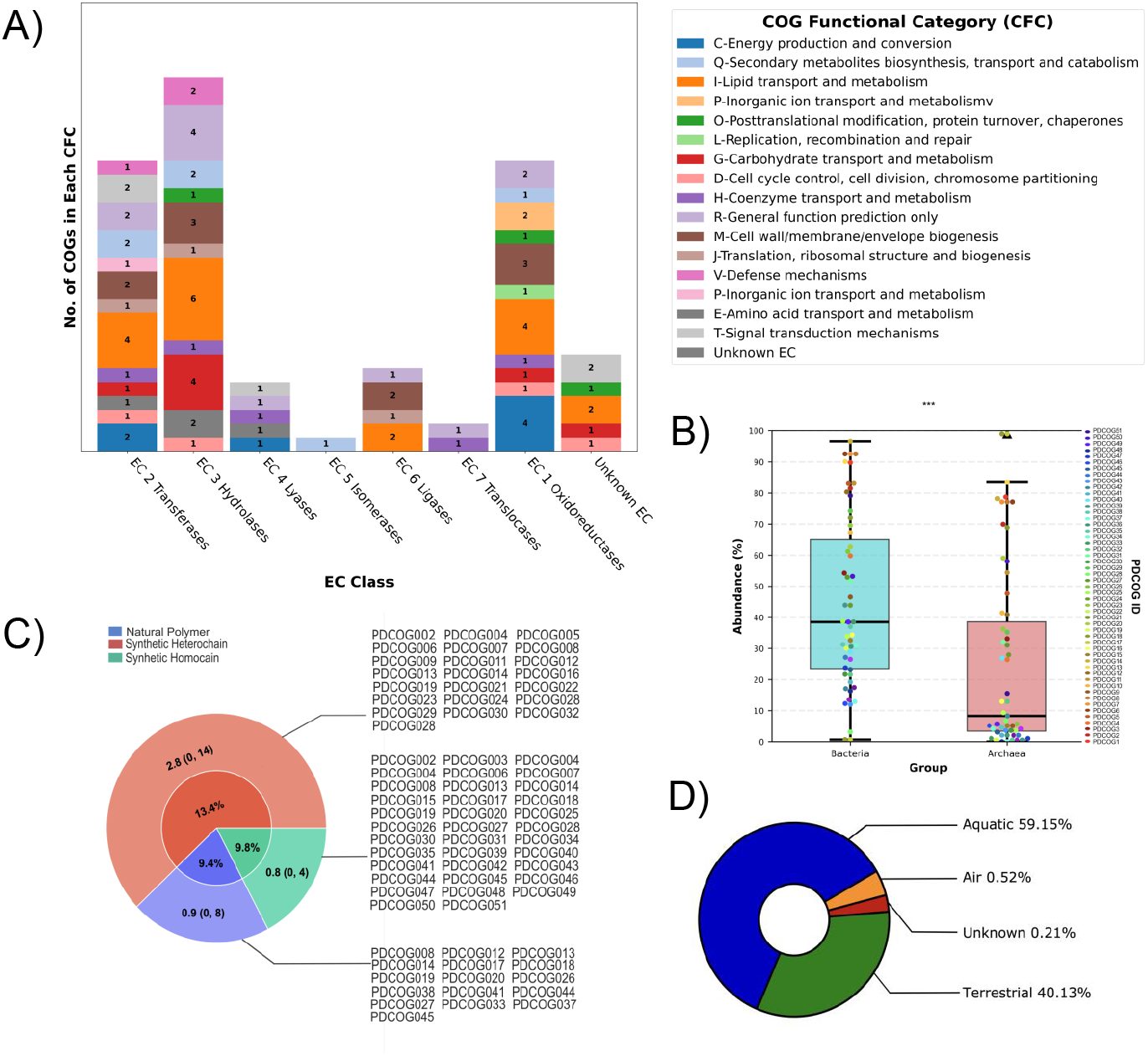
Functional, genomic and ecological diversity in the plastic-degrading clusters of orthologous groups (PDCOGs). A. Enzymatic roles and genomic functions of plastic biodegradation. The stacked bar plot illustrates the distribution of PDCOGs across COG functional categories (CFCs) within each EC class. The x-axis represents the EC classes (EC1 to EC7), which categorize enzymes on the basis of their catalytic reactions. Each bar is divided into colored segments, with each color representing a distinct CFC (e.g., metabolism, cellular processes, information storage). The count of correlated PDCOGs in each CFC is annotated within the corresponding segment of the bar, the sum of which, on the y-axis, indicates the quantification of COGs associated with each EC class. **B. Relative prokaryotic abundance in the PDCOGs**. Box plots showing the percentages of prokaryotes, bacteria, and Archaea (primary y-axis) across 51 PDCOGs (secondary y-axis), with a swarm plot overlay indicating enzyme-specific contributions (enzymes color-coded). The statistical significance of the dominance of bacteria over Archaea within prokaryotes was determined by a Wilcoxon signed-rank test (***: p < 0.001). The outliers are denoted by black triangles. **C. Plastic polymer biodegradation profile**. The pie chart shows the proportional distributions of natural, synthetic homochain, and synthetic heterochain plastic polymers potentially biodegradable by PDCOGs. The inner segment proportions indicate the relative degradation capacity of different categories of polymers, whereas the outer layer represents the average degradability with minimum and maximum counts. **D. Environmental distribution of PDCOGs**. Illustration of the mean proportional incidence of plastic degraders in various environmental habitats on the basis of genome counts from the Genome portal.

To assess enzyme-polymer specificity, we computed indicator values (IVs) for 212 EC subclasses across 39 polymers. While most associations had low IVs (75%, < 0.25), enzymes such as carboxylic ester hydrolases and glycosidases were broadly linked to many polymers. Notably, synthetic heterochain polymers were associated with more EC subclasses (n = 52) than were natural (n = 34) or synthetic homochain (n = 26) polymers, indicating greater enzymatic diversity for structurally complex plastics (Fig. 3B, Supplementary Dataset SD4).

**Fig. 3:**
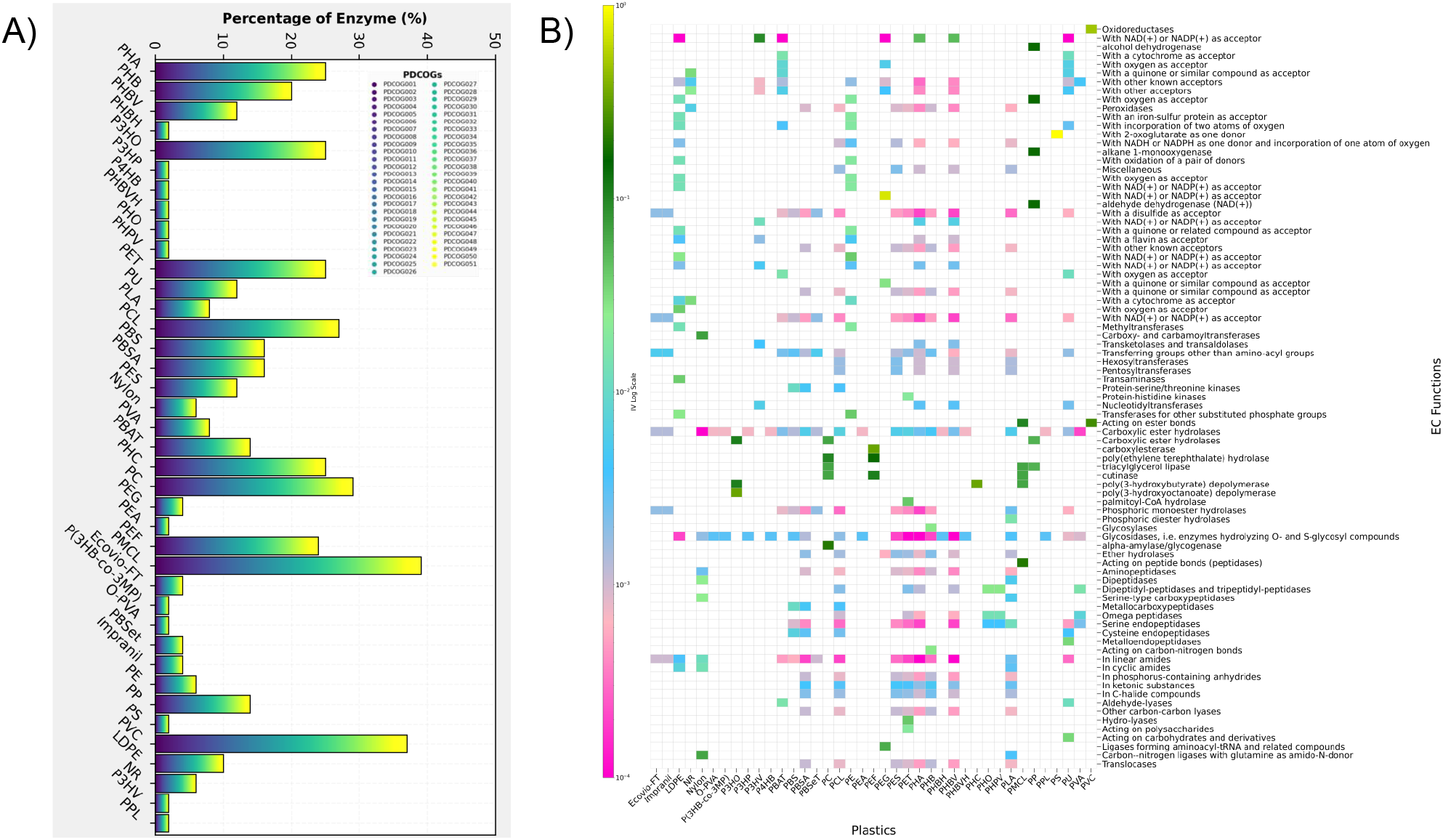
Enzymatic characterization of plastic-degrading clusters of orthologous groups (PDCOGs) on the basis of known biological enzymatic activities. A. Enzymatic activity of PDCOGs across plastic types. The bars represent the percentage of total enzyme activity (0–100%) among 51 PDCOGs for each plastic type. The x-axis displays plastic types, whereas the y-axis shows enzyme activity (%). Owing to the highest percentage being less than 50%, the y-axis only depicts up to 50% for better visualization. The legend (right) maps PDCOG domains to colors, with a purple-to-yellow gradient (bottom to top) indicating different protein clusters. **B. Heatmap of the indicator values (IVs) for enzymes associated with plastic biodegradation**. This plot shows the IVs for enzymes putatively involved in the biodegradation of 39 plastic types. With plastics on the x-axis and potential enzymes on the y-axis, the IV metric of relative abundance and specificity was calculated on the basis of annotations from the KEGG database ^44^. The color intensity and log-based value (10^−4^ lowest to 10^−0^ highest) highlight the strength of their association.

### Distribution of the PDCOGs across prokaryotes (archaea and bacteria)

Across 2,296 prokaryotic species (2,103 bacteria and 193 archaea), 13.4% were associated with synthetic heterochain polymers, 9.8% with synthetic homochain polymers, and 9.4% with natural polymers (Fig. 2C; Table 1). Bacteria displayed a significantly broader distribution of PDCOGs than archaea did (the median PDCOGs in bacteria was 38.5%, and the median PDCOGs in archaea was 8.3%, with an overall prevalence of 35.4% in prokaryotes; Wilcoxon signed-rank test, p < 0.0001) (Fig. 2B, Supplementary dataset SD1).

On average, the bacterial and archaeal genomes encode 20 and 11 PDCOGs (out of 51), respectively (Supplementary dataset SD1). The largest PDCOGs in archaea were found in Asgardarchaeota (n = 17) and Euryarchaeota (n = 15), whereas Myxococcota (n = 38) and Acidobacteriota (n = 29) were the largest groups in bacteria. Overall, 96.6% of prokaryotes have at least one PDCOG, i.e., a copy of a PPDP. PDCOG015, a chaperonin cofactor present in 99.0% of archaeal species, has protease activity with the potential to breakdown PLA ^28^. In bacteria, PDCOG012, which has dehydrogenase activity, has the potential to participate in the intracellular biodegradation of P3HV, PHBV, and PHA ^28^. In contrast, PDCOG034 (orthologous group with chitinase activity) in archaea and PDCOG015 in bacteria are poorly represented across the respective lineages.

### Phyletic distribution of the PDCOGs in natural environments

To evaluate the taxonomic overlap in degradation potential, we used BLAST-derived top hits from seed sequences to assess the shared microbial capacities across plastic types (Fig. 4A). We analyzed the relative abundance of the resulting species to determine which plastic types have the most diverse biodegradability potential (synthetic heterochain > natural > synthetic homochain) and which taxa are most frequently associated with specific plastics (Pseudomonadata, Actinomycetota, Bacillota, and Bacteroidota) (Fig. 4B). Synthetic heterochain polymers harbored the highest proportion of unique microbial degraders (88%), followed by natural (65%) and synthetic homochain (32%) polymers. Notably, some strains have the potential capacity to biodegrade more than one polymer type. Only 22% of the species were shared across all three categories, with the greatest overlap between natural and synthetic heterochain polymers (41%); no species were shared between synthetic homochain polymers and the other groups (Fig. 3A).

**Fig. 4.**
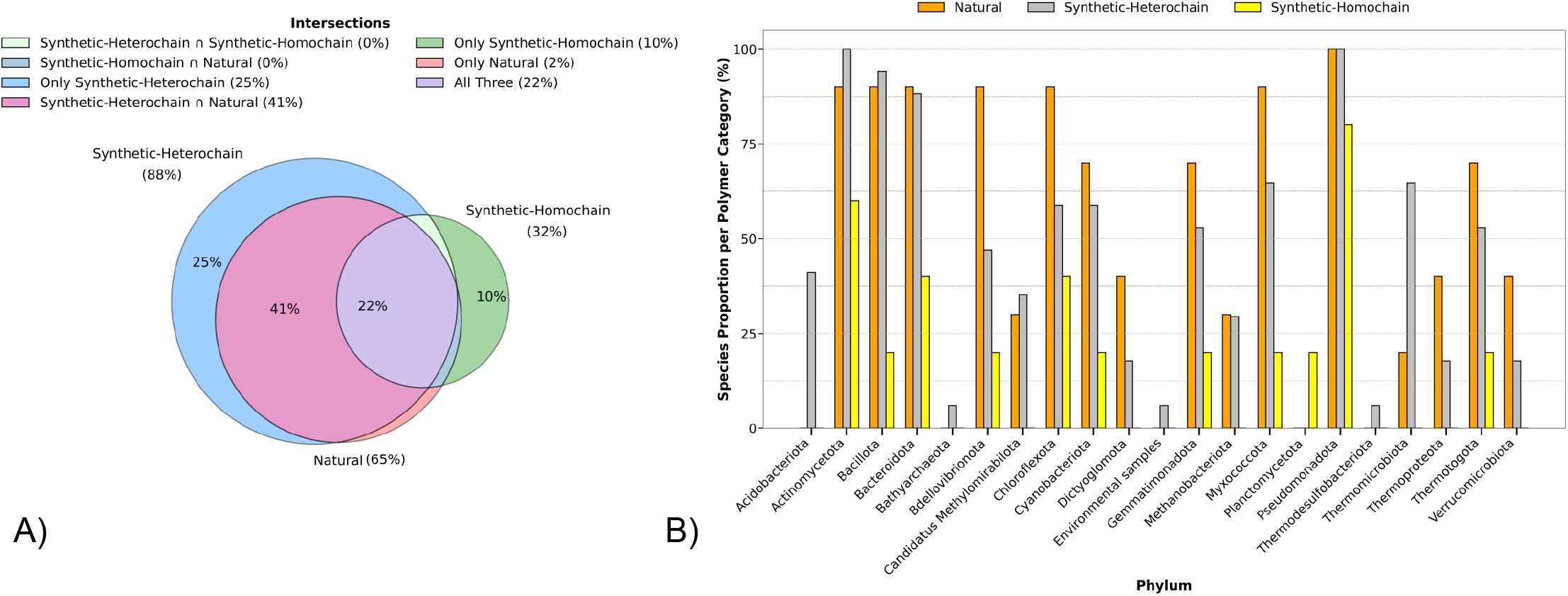
Taxonomic distribution of plastic-degrading microbes and polymer types. A. Species-level distribution patterns among polymer classes. Venn diagram illustrating the relative distributions of common species among the plastic polymer categories natural, synthetic heterochain, and synthetic homochain for biodegradation. Each circle represents one plastic category with color grading, and the intersections denote shared associations among multiple categories. Percent values are shown for major intersections, whereas low-frequency overlaps (<5%) are denoted in the legend for clarity. Circle positions and sizes vary on the basis of their affinity with species. **B. Phylum-specific trends in categorical polymer biodegradation**. The grouped bar plot compares the relative associations of phyla within each plastic category. Each bar cluster on the x-axis corresponds to all the unique prokaryotic phyla in the dataset. The height of each bar along the y-axis denotes the phylum-based proportion of unique species that are correlated with the plastics in each category (natural, synthetic heterochain, synthetic homochain), normalized by the total unique plastics within that category.

Most prokaryotic species present enzymes with the capacity to break down plastic polymers (Fig. 4B). PDCOGs are widely distributed across prokaryotic species (Fig. 1, Supplementary Fig. S1, Supplementary Dataset SD3). In nature, Pseudomonadota is the bacterial group that presents the greatest abundance of PPDP of all polymer types (natural and synthetic), followed by Actinomycetota, Bacteroidota and Bacillota (Fig. 4B). In archaea, three phyla, namely, Methanobacteriota, Thermoproteota and Bathyarchaeota, were identified as potential biodegraders of plastic polymers. Our analysis revealed that 65.68% of the cataloged species potentially biodegrade natural polymers, 32.47% of the species biodegrade synthetic homochain polymers, and 87.82% biodegrade synthetic heterochain polymers.

### Environmental and geographic distributions of the PDCOGs

PPDPs have been detected across a broad spectrum of aquatic and terrestrial environments (Fig. 5; Table 1). Although aquatic environments dominated the dataset due to sampling bias, several PDCOGs (e.g., PDCOG034, PDCOG035, PDCOG045, and PDCOG050) were enriched in continental samples. Geographic data indicate that terrestrial sequences are derived primarily from North America, Western Europe, East Asia, and Australia, whereas marine sequences reflect typical oceanographic sampling routes. Statistical comparison with the expected distributions (chi-square test; Supplementary Table S3) revealed a general underrepresentation of PPDPs in the environmental samples, with the exception of endolithic (rock-dwelling) niches, which were significantly enriched. The soil samples presented the expected representation, but the PDCOG frequencies varied by habitat (Supplementary Dataset SD6), suggesting environment-specific selective pressures on plastic-degrading functions.

**Fig. 5.**
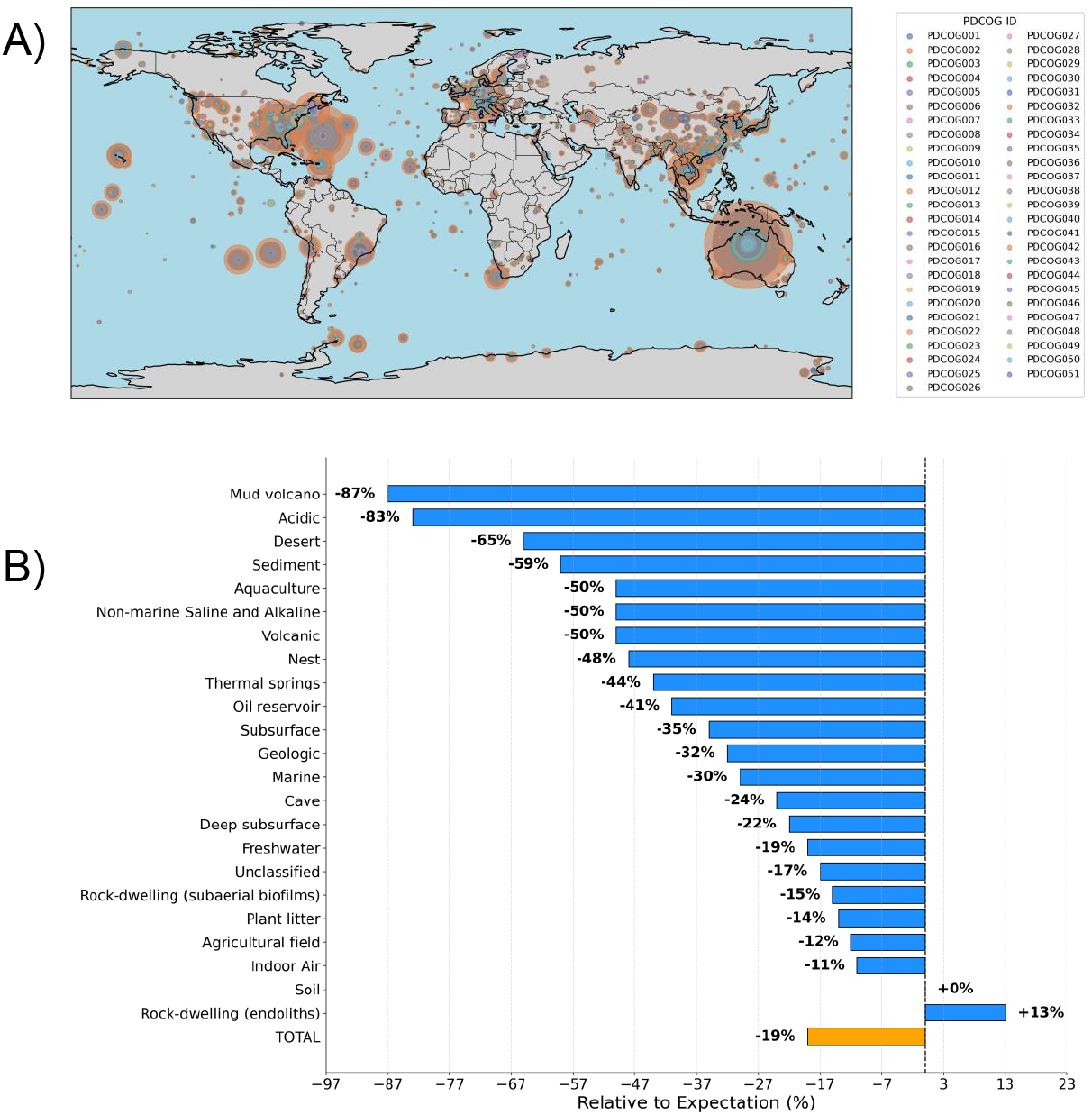
Global distribution and abundance of plastic-degrading proteins. A. World distribution of plastic-degrading clusters of orthologous groups (PDCOGs) across environments. The geospatial distribution of PDCOGs highlights hotspots of functional diversity and the frequency of plastic-degrading genes across the sampled environments. The color gradients represent functional diversity of PDCOGs, whereas their proportional variation denotes sequence abundance. Hotspots of enzymatic potential correlate with anthropogenic impact zones (e.g., soil, sediment, marine and aquatic) and microbial niches adapted to synthetic polymer hydrolysis. **B. Percentages of observed sequences relative to expectations (O/E) across various environments**. The observed (O) and expected values (E) were calculated on the basis of the data obtained from the JGI Genome Portal and the NCBI COG database, respectively. The values represent the percentage of deviation from the expected baseline (100%). The bars to the left of the central axis indicate environments where the observed values fall below expectations, whereas the bars to the right represent environments exceeding expectations. The “TOTAL” summary is displayed in orange for distinction.

## DISCUSSION

### General characteristics of the PDCOGs

The PDCOG database provides a comprehensive and functionally annotated collection of PPDPs organized into orthologous groups on the basis of sequence homology and enzymatic activity. These proteins span diverse COG functional categories, with hydrolases (EC3s) being the most prevalent, followed by oxidoreductases (EC1s) and transferases (EC2s). The most represented COG functional categories associated with plastic degradation are lipid transport and metabolism (Category I), cell wall/membrane/envelope biogenesis (Category M), and energy production and conversion (Category C). This diversity reflects the biochemical complexity of plastic degradation, which often requires multistep enzymatic cascades. Notably, all proteins in PDCOGs are derived from sequences with known or predicted enzymatic functions. However, we hypothesize that ongoing evolutionary pressures, particularly in plastic-contaminated environments, may drive the emergence of novel PPDPs. These could arise through mechanisms including horizontal gene transfer, lineage-specific gene loss, and rapid sequence diversification, potentially resulting in enzymes with minimal similarity to previously characterized biodegradative proteins ^29^.

Differences in polymer structure appear to significantly influence biofilm formation by prokaryotes and thus the distribution of PDCOGs. Overall, synthetic heterochain polymers were associated with a greater number of unique PPDP-hosting species (25%) than were synthetic homochain (10%) and natural polymers (2%). This analysis is based on seed sequences ^28^ derived from prokaryotic taxa isolated from either natural environments or laboratory experiments, all of which demonstrate some capacity for polymer biodegradation. However, the presence of a PPDP does not necessarily equate to enzymatic activity *in situ*. This discrepancy may be due to three principal factors: (1) incomplete sampling of microbial diversity, (2) the potential for enzymatic inactivity in nonnative or suboptimal environmental conditions, and (3) putative inactivation of the enzyme’s main functional domain. Additionally, structural alterations of plastics in natural environments—via photodegradation, oxidation, or mechanical stress —are not fully understood and may influence microbial colonization and enzymatic access.

### Phyletic distribution of the PDCOGs

Since the term *plastisphere* was first introduced in 2013 to describe microbial communities associated with plastic debris in marine environments, considerable progress has been made in understanding the phylogenetic and functional diversity of plastic-degrading microbes ^30,31^. Here, we show that bacteria encode a broader repertoire of plastic-degrading enzymes than archaea do across the surveyed environments. This dominance likely reflects both the greater numerical abundance of bacteria and their greater metabolic versatility, enabling colonization across diverse plastisphere niches. Nevertheless, archaea also harbor functionally distinct PDCOGs, suggesting that niche-specific or extremophile adaptations remain underexposed.

Strikingly, over 95% of the surveyed prokaryotic species possess at least one PDCOG, including >98% of archaea and >96% of bacteria; thus, these species have the potential to biodegrade at least one polymer type. Among bacteria, *Pseudomonas sp*. has the greatest number of PPDPs found in natural environments, which is consistent with its known metabolic versatility and adaptability^32^. For example, clinical isolates of *Pseudomonas aeruginosa* have been shown to be able to degrade medical-grade plastics^33^, whereas wild-type strains are efficient at remediating petrochemical spills^34^. Other prominent bacterial groups enriched in PPDPs include Actinomycetota, Bacteroidota, and Bacillota, which aligns with prior analyses of 436 putative plastic-degrading taxa, where Proteobacteria (37.1%), Ascomycota (16.9%), Actinobacteria (14.3%), Firmicutes (13.8%), and Basidiomycota (8.1%) were among the most represented phyla^35^. We identified three archaeal phyla—Methanobacteriota, Thermoproteota, and Bathyarchaeota—as potential plastic degraders, underscoring a likely underappreciated role for archaea in plastic biodegradation, particularly in extreme or anaerobic environments ^35^.

### Geographical and environmental distributions of the PDCOGs

Although PDCOGs are most commonly identified in aquatic systems, terrestrial samples—particularly soil and lithic (rock-associated) environments—also present disproportionately high levels of PPDPs. We propose that nutrient limitations in these terrestrial habitats may exert selective pressure on the acquisition or maintenance of plastic-degrading enzymes^36^. In contrast, marine environments—despite their relative nutrient scarcity—benefit from the accumulation of dissolved organic carbon, nitrogen, and phosphorus, potentially supporting broader microbial functionality ^37^.

It is important to acknowledge potential limitations in genome representation across databases, which may contribute to underestimations of PDCOG abundance in certain environments. Nonetheless, our findings emphasize the importance of environmental specificity in shaping the distribution of plastic-degrading functions and suggest that microbial adaptation to plastic pollution may be ongoing and context dependent.

### Applications and future directions

The PDCOGs database integrates functional, ecological, and taxonomic information, providing a critical reference for researchers studying microbial plastic degradation. It complements existing resources on plastic-degrading proteins and microbes ^28,38–41^. Biodegradable polymers, such as polycaprolactone (PCL), polybutylene succinate-coadipate (PBSA), polybutylene succinate (PBS), and polylactic acid (PLA), are more amenable to enzymatic degradation because of their ester linkages^42,43^. In contrast, high-molecular-weight polymers such as PET require highly efficient and specific enzymes for depolymerization. PDCOGs provide a framework to explore such structure-function relationships and guide the identification of candidate enzymes for biotechnological and material science applications.

Addressing the global plastic pollution crisis, particularly the proliferation of MNPs, will require integrative, cross-disciplinary strategies combining genomics, biochemical characterization, synthetic biology, biochemical engineering and environmental microbiology. Given that biodegradation efficacy is strongly influenced by both environmental conditions and polymer structure, PDCOGs provide a foundational resource for advancing our understanding of microbial plastic degradation potential across ecosystems.

## METHODS

### Seeds of putative plastic-degrading proteins

A seed of putative plastic-degrading protein (PPDP) sequences was obtained from the PlasticDB ^28^ and articles from the literature to expand the coverage of the dataset (Supplementary Table S4). The initial dataset (available from the PDCOGs webpage) contains a seed of PPDP sequences from bacterial species and potential plastic polymers that can be biodegraded (Supplementary Table S5).

### Orthologous proteins

We collected all proteins from the COGs database ^43^. This database contains 5,640,669 protein sequences classified on the basis of orthologous relationships into 5,050 COGs. Each protein is functionally annotated, and the COGs are classified into 26 biological functions on the basis of four main functional categories (Supplementary Table S2). As of the 2024 update, the COGDB includes 2,296 pieces of genomic information (2,103 bacterial and 193 archaeal).

### Putative plastic-degrading orthologous groups and phyletic patterns

The PDCOG dataset was created on the basis of our BLASTp results from the NCBI COG database (COGDB) ^43^ and PlasticDB ^28^ (Supplementary Datasets SD1-3). Each sequence from the PPDP dataset was mapped onto the COG dataset with the program BLASP for the identification of potential plastic-degrading orthologous groups. Briefly, a BLASTP search was performed on the NCBI COG database and the PlasticDB database to obtain the best COG hits against the PlasticDB (BLASP parameters were outfmt = 6, *e*-value = 0.001, max-target-seqs = 5). The COG functional category and symbol data were collected from the COGDB.

### Functional annotation of the PDCOGs

The functional annotations of the PDCOGs (Supplementary Dataset SD4) were performed with the database Cluster of Orthologous Group Database (COGDB) ^43^, Kyoto Encyclopedia of Genes and Genomes (KEGG) ^44^, InterPro 104.0 ^45^, ExPASy ^46^, and NCBI Taxonomy ^47^. The KEGG database was used to collect KO codes for each PDCOG, as well as the EC (enzymatic) and reaction codes. The categorical function of each EC class was characterized with ExPASy. The protein functional domain and gene ontology (GO) information of each PDCOG was characterized with the InterPro database.

### Reconstruction of the polymers dataset

The systematic workflow for the data curation of the polymers included in the dataset is based on both cheminformatics analyses and literature reviews (Supplementary Table S5 and Supplementary Dataset SD2). We used the PU-REST API of Pubchem DB ^48^ for SMILEs, the Polymer Genome DB, Smiles2Monomers ^49^ for monomer mapping, RDKit open source cheminformatics v2025.3.2 in Python for validation of SMILES and monomers, and Beautiful Soup for scraping open access journals from PubMed Central ^50^. Among the 39 polymers, 12 PSMILEs were obtained from PubChem and the rest were obtained from PI1M DB ^51^ for full repeat units vs. fragments, using the filters ‘™Atomcount ≥ 10 to exclude fragments, ™Molecular weight 500–5000 Da, Atom count, functional groups, and excluding Non-specific’. Consequently, after the 50 best batches per target polymer were obtained, consensus-based PSMILEs were obtained for the remaining polymers. To evaluate recent advances in the potential biodegradation of additional polymers, including P3HO, PC, PEF, PMCL, PP and PPL, we surveyed 300 peer-reviewed studies published between 2000 and 2025. The selection specifically relied on prokaryotic relevance to these polymers and enzymes used in biodegradation.

### Environmental annotation of the PDCOGs

Protein sequences of free-living species with geographical and environmental information were obtained from the Joint Genome Institute (DOE-JGI) ^52^. We analyzed 74,651,773 protein sequences with the Python *re* package to identify sequences with geographical and environmental annotations that belong to free-living microbial species. The filtered sequences were mapped on the PDCOGs with BLASTP. Thus, the final dataset included 662,830 protein sequences with environmental and functional annotations (Supplementary Table S3 and Supplementary Datasets SD5-6). These proteins are spread across the Earth in 23 different environments. The distribution map of the PDCOGs was built using pandas, matplotlib, cartopy.crs for handling world map projections, and seaborn for the built-in color palettes. A chi-square test was performed on the PDCOGs dataset to determine significant differences (p<0.05) from an expected distribution of the protein sequences on the basis of protein number in the COGDB.

### Plotting and statistical analysis

All data processing, statistical analyses and visualizations were performed with Python (version 3.9.6). Visualizations were generated using Matplotlib (v3.9.4), Plotly (v6.0.0.) and Seaborn (v0.13.2), whereas data processing was performed with the Pandas (v2.2.3), SciPy (v1.13.1), and NumPy (v2.0.2) libraries and the scipy.stats module (wilcoxon). The Wilcoxon signed-rank test (Figure 2B) was used to assess significant differences (p value < 0.05) in the PDCOG abundance distribution between the bacteria and archaea. The indicator value (IV) in the heatmap (Fig. 3B) was calculated. Briefly, the method combines specificity (Sp) and fidelity (Fd), which are calculated for each enzyme-plastic pair ^53^, and downweighting is applied to reduce the influence of frequently occurring enzymes (Equation 1). A permutation test (10,000 iterations) was used to assess significance, yielding p values ranging from 0.01 to 0.04 (Fig. 2B).

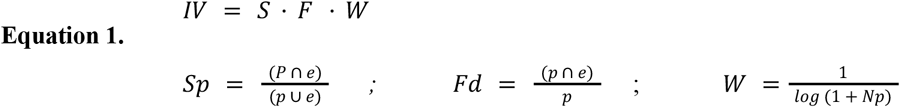

Where p is the total EC occurrences for the target plastic polymer; e is the total 4th-subclass EC occurrences under the 3rd-subclass EC across plastics; p⋂e is the number of EC occurrences (4th EC subclass) for the target plastic under the 3rd EC subclass; p⋃e is the union of the 3rd EC subclass annotation of the target plastic and across plastics; and W is a downweighting factor.

## AUTHOR CONTRIBUTIONS

Conception and design of the study: MN, PP; Funding acquisition: MN, KS, PP; Data collection: SM, LTF, PP; Data analysis: SM, LTF, PP; Manuscript drafting: SM, LTF, PP.; Manuscript revision for critical intellectual content: SM, LTF, KS, MN, PP; Writing the final version of the manuscript: MN, PP. All authors have read and agreed to the published version of the manuscript.

## ACKNOWLEDGEMENTS

We thank members of the MicroWorld team, students and collaborators for their helpful discussions.

