## Supplementary Material for "Plastic-degrading clusters of orthologous groups reveal near-universal biodegradation potential in prokaryotes" (from another version of this preprint: version 1, 2025-06-10)

### SUPPLEMENTARY FIGURES

**Supplementary Figure S1.** Bipartite networks of plastic-degrading microbes, orthologous groups and polymers. Full data available at https://phylobone.com/microworld/PDCOG.

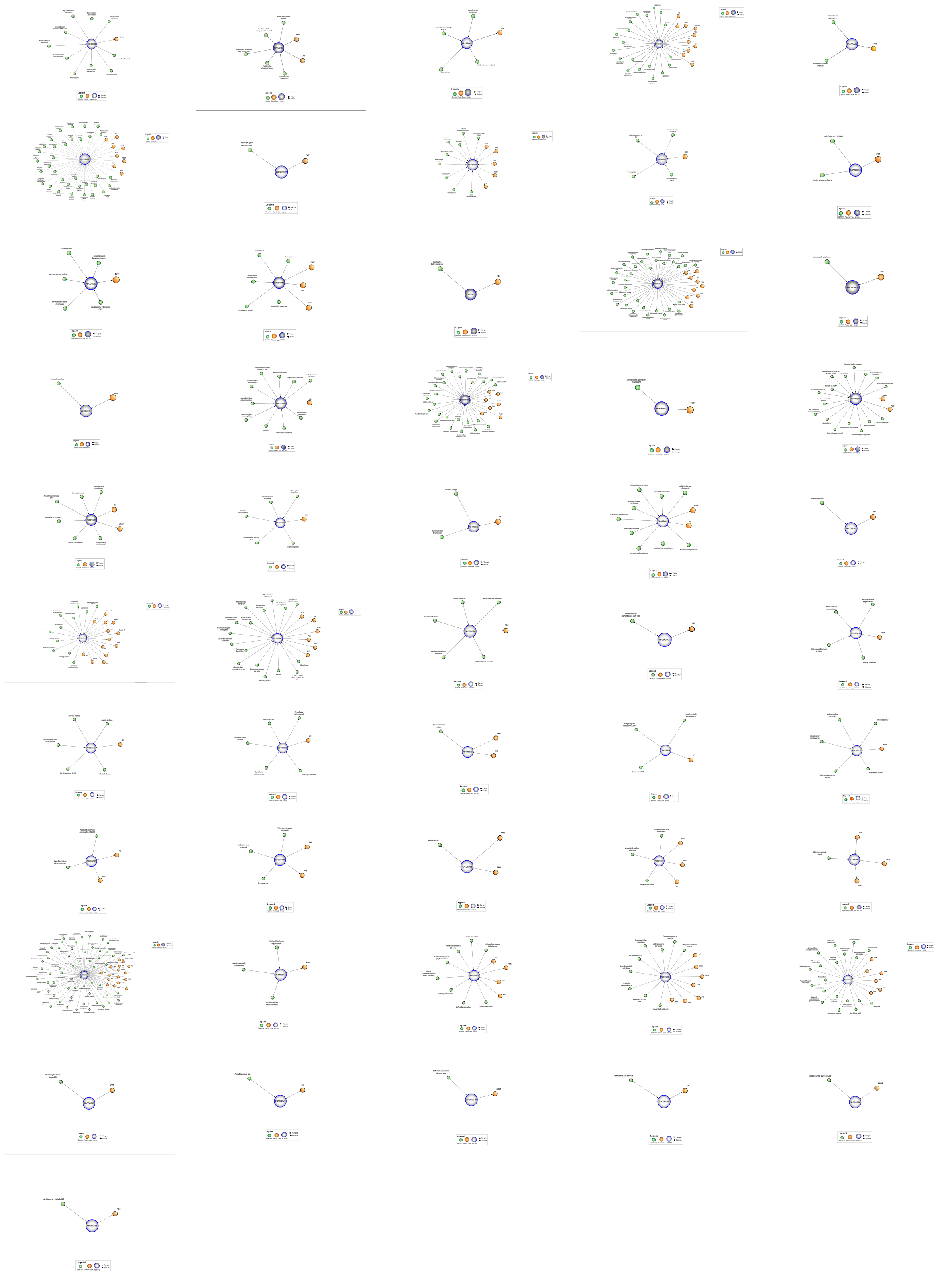

**Supplementary Figure S2.** Environmental distribution of plastic-degrading COGs (PDCOGs). Distributions of PDCOGs on the world map. The map was plotted using the metadata of each PDCOG, with colors and size indicating the number of the sequences found at that location. Full data freely available at [https://phylobone.com/microworld/PDCOG](https://phylobone.com/microworld/PDCOG).

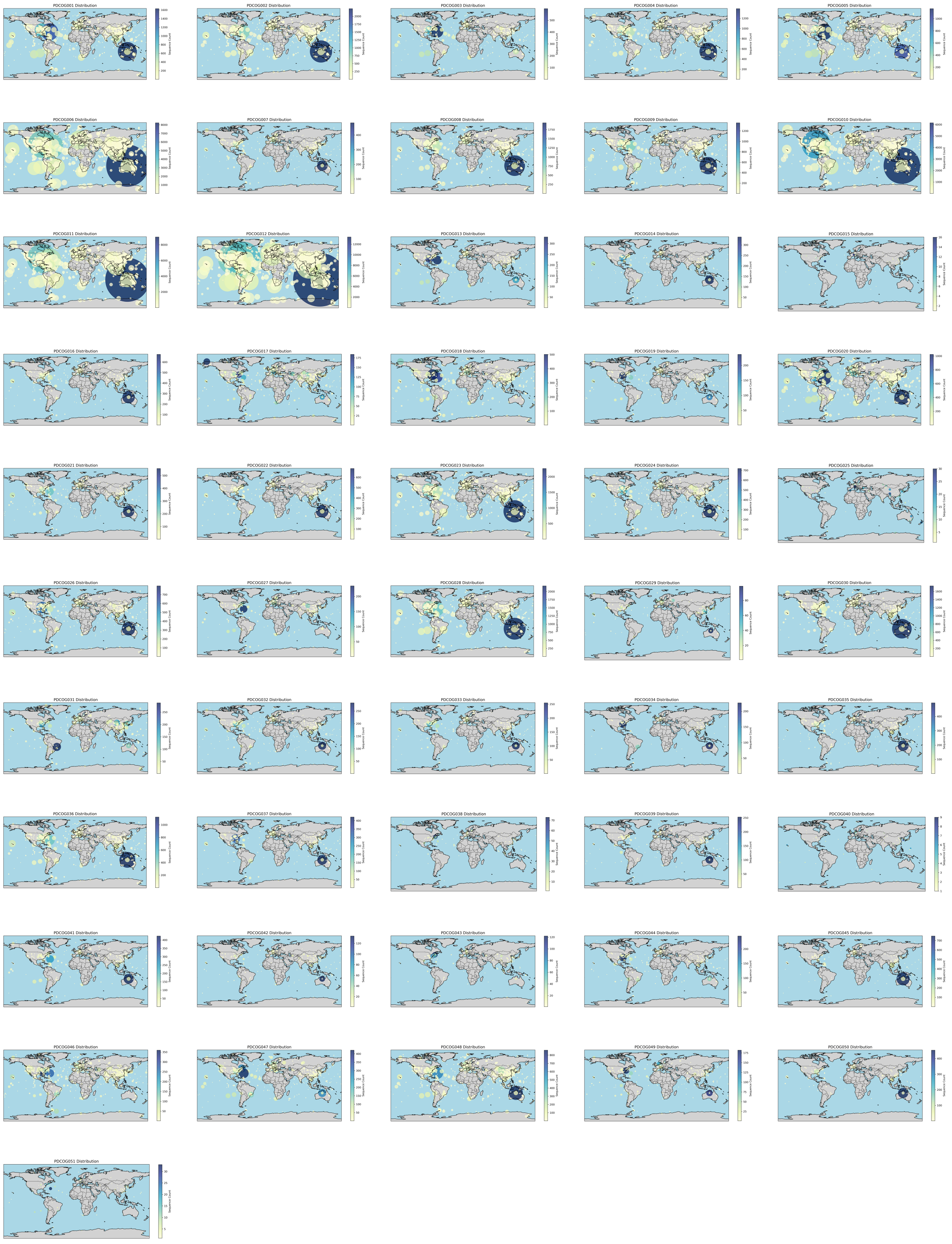

**SUPPLEMENTARY TABLES**

**Supplementary Table S1.** Resources available for plastic-degrading proteins

| Resource | Protein sequences | Orthologous groups | Microbes | Phyletic Patterns | Enzyme information | Functional characterization | Protein domains | Geographical location (lat,long) | Environmental information | Ref. |
|---|---|---|---|---|---|---|---|---|---|---|
| PDCOGs | Yes (n = 662,830) | Yes (n = 51) | Yes | Yes | Yes | Yes | Yes | Yes | Yes | - |
| PlasticDB | Yes (n =228) | | Yes | Yes | Yes | | | | | [31] |
| PAZy | Yes (n =256) | | Yes | | Yes | Yes | | | | [32] |
| PMDB | Not available | Yes | Yes | Yes | | | | | Yes | [33] |
| MIBPOP | Yes (n = 5,736) | | Yes | Yes | Yes | Yes | | | | [34] |
| UM-BBD/EAWAG-BBD | This website is no longer actively maintained. | | | | | | | | | [35] |

**Supplementary Table S2.** Functional categories included in the database of clusters of orthologous groups (COGs) [30]

| INFORMATION STORAGE AND PROCESSING | | CELLULAR PROCESSES AND SIGNALING | | METABOLISM | | POORLY CHARACTERIZED | |
|---|---|---|---|---|---|---|---|
| **J** | Translation, ribosomal structure and biogenesis | **D** | Cell cycle control, cell division, chromosome partitioning | **C** | Energy production and conversion | **R** | General function prediction only |
| **A** | RNA processing and modification | **Y** | Nuclear structure | **G** | Carbohydrate transport and metabolism | **S** | Function unknown |
| **K** | Transcription | **V** | Defense mechanisms | **E** | Amino acid transport and metabolism | | |
| **L** | Replication, recombination and repair | **T** | Signal transduction mechanisms | **F** | Nucleotide transport and metabolism | | |
| **B** | Chromatin structure and dynamics | **M** | Cell wall/membrane/envelope biogenesis | **H** | Coenzyme transport and metabolism | | |
| | | **N** | Cell motility | **I** | Lipid transport and metabolism | | |
| | | **Z** | Cytoskeleton | **P** | Inorganic ion transport and metabolism | | |
| | | **W** | Extracellular structures | **Q** | Secondary metabolites biosynthesis, transport and catabolism | | |
| | | **U** | Intracellular trafficking, secretion, and vesicular transport | | | | |
| | | **O** | Posttranslational modification, protein turnover, chaperones | | | | |
| | | **X** | Mobilome: prophages, transposons | | | | |

**Supplementary Table S3.** Distribution of potential plastic-degrading proteins (PPDP) across 23 environments.

| Environment | (O) Number of PD-Proteins | Number of Proteins | Frequency (O) | (E) Expected PD-Proteins | Frequency (E) | (O/E)·100 | Chi square | p-Value | Result |
|---|---|---|---|---|---|---|---|---|---|
| Acidic | 96 | 15827 | 0.006 | 557 | 0.035208 | 17% | 381.8 | <0.001 | LESS |
| Agricultural field | 1277 | 41129 | 0.031 | 1448 | 0.035208 | 88% | 20.2 | <0.001 | LESS |
| Aquaculture | 265 | 15013 | 0.018 | 529 | 0.035208 | 50% | 131.4 | <0.001 | LESS |
| Cave | 877 | 32721 | 0.027 | 1152 | 0.035208 | 76% | 65.7 | <0.001 | LESS |
| Deep subsurface | 3521 | 128576 | 0.027 | 4527 | 0.035208 | 78% | 223.5 | <0.001 | LESS |
| Desert | 860 | 69820 | 0.012 | 2458 | 0.035208 | 35% | 1039.1 | <0.001 | LESS |
| Freshwater | 114514 | 4005335 | 0.029 | 141018 | 0.035208 | 81% | 4981.3 | <0.001 | LESS |
| Geologic | 2909 | 121811 | 0.024 | 4289 | 0.035208 | 68% | 443.8 | <0.001 | LESS |
| Indoor Air | 1191 | 37858 | 0.031 | 1333 | 0.035208 | 89% | 15.1 | <0.001 | LESS |
| Marine | 198575 | 8112561 | 0.024 | 285623 | 0.035208 | 70% | 26529.3 | <0.001 | LESS |
| Mud volcano | 48 | 10664 | 0.005 | 375 | 0.035208 | 13% | 285.6 | <0.001 | LESS |
| Nest | 869 | 47472 | 0.018 | 1671 | 0.035208 | 52% | 385.2 | <0.001 | LESS |
| Non-marine Saline and Alkaline | 5850 | 329962 | 0.018 | 11617 | 0.035208 | 50% | 2863.0 | <0.001 | LESS |
| Oil reservoir | 505 | 24516 | 0.021 | 863 | 0.035208 | 59% | 148.6 | <0.001 | LESS |
| Plant litter | 1063 | 35179 | 0.030 | 1239 | 0.035208 | 86% | 24.9 | <0.001 | LESS |
| **Rock-dwelling (endoliths)** | **1100** | **27593** | **0.040** | **971** | **0.035208** | **113%** | **17.0** | **<0.001** | **MORE** |
| Rock-dwelling (subaerial biofilms) | 248 | 8279 | 0.030 | 291 | 0.035208 | 85% | 6.5 | 0.01 | LESS |
| Sediment | 10336 | 723640 | 0.014 | 25478 | 0.035208 | 41% | 8998.8 | <0.001 | LESS |
| **Soil** | **261948** | **7465491** | **0.035** | **262841** | **0.035208** | **100%** | **3.0** | **0.08** | **EXPECTED** |
| Subsurface | 74 | 3249 | 0.023 | 114 | 0.035208 | 65% | 14.3 | <0.001 | LESS |
| Thermal springs | 6595 | 335040 | 0.020 | 11796 | 0.035208 | 56% | 2293.1 | <0.001 | LESS |
| Unclassified | 49651 | 1702948 | 0.029 | 59957 | 0.035208 | 83% | 1771.4 | <0.001 | LESS |
| Volcanic | 458 | 25920 | 0.018 | 913 | 0.035208 | 50% | 226.4 | <0.001 | LESS |
| TOTAL | 662830 | 23320604 | 0.028 | 821061 | 0.035208 | 81% | 50868.9 | <0.001 | LESS |

***Frequency (E) = N_PDCOG_Proteins / N_Proteins_COG_Database =*** *198595 / 5640699 = 0.035208*

**Supplementary Table S4.** List of articles used to identify additional plastic-degrading enzymes.

| Polymer | Species | TAXID | Putative Enzyme | Uniprot ID | Functional domain / Functional Class | EC Number | Ref. |
|---|---|---|---|---|---|---|---|
| P3HO | *Alcaligenes faecalis AE122* | 511 | Poly(3-hydroxybutyrate) depolymerase (PHB depolymerase) | P94146 | Not specified | Not specified | [1] |
| | *Pseudomonas lemoigne* | 29443 | Poly(3-hydroxybutyrate) depolymerase (PHB depolymerase) | not found | Not specified | Not specified | [2] |
| | *Pseudomonas putida* | 160488 | PHA depolymerases | Q9I6R6 | Alpha/beta hydrolase fold | 3.1.1.75 | [3] |
| | *Cupriavidus necator* | 381666 | PHA depolymerase | Q0K9J1 | Ser-His-Asp triad | 3.1.1.75 | [4] |
| | *Streptomyces exfoliatus* | 1916 | PHA depolymerase | P71084 | Lipase-like | 3.1.1.75 | [5] |
| | *Pseudomonas fluorescens GK13* | 294 | Polyhydroxyalkanoate depolymerase, polyhydroxyalkanoate (MCL-PHA) | Q3L8T3 | Hydrolase | 3.1.1.76 | [6] |
| | *Rhodospirillum rubrum* | 1085 | PHA depolymerase | Q2RQN1 | PHA_depol_C | 3.1.1.75 | |
| PHC | *Pseudomonadaceae* | 135621 | PHA depolymerases | Not specified | Not specified | 3.1.1.75<br>3.1.1.76 | [7] |
| | *Cupriavidus necator* | 106590 | | | | | |
| | *Arenicella spp.* | 1872634 | PHA depolymerase | Not specified | Not specified | Not specified | [8] |
| | *Microbacterium spp.* | 51671 | | | | | |
| PVC | *Pseudomonas citronellolis* | 29461 | PVC-associated dehalogenase | Q9I6R6 | Haloacid dehalogenase | 3.8.1.5 | [9] |
| | *Bacillus flexus* | 86664 | Not specified | Not specified | Not specified | Not specified | |
| | *Bacillus licheniformis* | 1402 | Not specified | Not specified | Not specified | Not specified | [10] |
| | *Achromobacter xylosoxidans* | 85698 | Not specified | Not specified | Not specified | Not specified | |
| | *Citrobacter koseri* | 545 | Not specified | Not specified | Not specified | Not specified | [11] |
| | *Klebsiella sp. EMBL-1* | 244366 | Catalase-peroxidase | Not specified | Peroxidase domain | 1.11.1.21 | |
| | | | Hydrolases (acting on ester bonds) | Not specified | Hydrolases (acting on ester bonds) | EC 3.1.-.- | [12] |
| | | | Oxidoreductases involved in various oxidative processes | Not specified | Oxidoreductases | EC 1.-.-.- | |
| PC | *Pseudoxanthomonas sp. strain NyZ600* | 1871049 | Hydrolase activity | Not specified | Hydrolase | 3.-.-.- | [13] |
| | *Bacillus subtilis* | 1423 | Cutinase | Q47QG0 | Alpha/beta hydrolase | 3.1.1.74 | [14] |
| | *Bacillus megaterium* | 1404 | Amylase, Lipase | Not specified | Not specified | 3.1.1.3 | [15] |
| | *Pseudomonas sp.* | 306 | Esterases, Lipase | P20845, A0AA86LWP4 | Hydrolases (acting on ester bonds) | EC 3.2.1.1 | [16] |
| PP | *Dermacoccus sp. strain AYDL3* | 1423 | Lipase (EstB) | O07553 | Alpha/beta hydrolase | 3.1.1.3 | [17] |

| | | | | | | | |
|---|---|---|---|---|---|---|---|
| | *Nocardioides sp.* | 35761 | Alkane hydroxylase | Not specified | Not specified | Not specified | 18 |
| | *Arenimonas sp.* | 1872635 | Alcohol dehydrogenase (Hartmannibacter) | Not specified | Oxidoreductases (Alcohol metabolism) | EC 1.2.1.3 | |
| | *Bradyrhizobium sp.* | 376 | copper oxidase | Not specified | Not specified | Not specified | |
| | *Pseudomonas sp.* | 306 | Laccase | Not specified | Not specified | 1.10.3.2 | |
| | *Bacillus sp. strain 27* | 519738 | Not specified | Not specified | Not specified | Not specified | 19 |
| | *Rhodococcus sp. strain 36* | 1831 | Not specified | Not specified | Not specified | Not specified | |
| PEF | *Ideonella sakaiensis* | 1547932 | PET hydrolase | A0A0K8P6T7 | Lipase | 3.1.1.101 | 20 |
| | *Marine biodegradation* | N/A | Not specified | Not specified | Not specified | Not specified | 21 |
| PMCL | *Pseudomonas fluorescens* | 294 | PHA depolymerase | Q9F5G3 | Lipase-like | 3.1.1.75 | |
| | *Alteromonas sp. Ghpt-2* | 232 | Lipase | Not specified | Lipase | 3.1.1.3 | 22 |
| | | | Cutinase | | Cutinase | 3.1.1.74 | |
| | *Ectopseudomonas hydrolytica DSWY01T* | | Cutinase | | | 3.1.1.74 | 23 |
| | | 2493633 | Lipase | Not specified | Lipase | 3.1.1.3 | |
| | *Fusarium solani* | 169385 | Cutinase (Rcut) | Not specified | Cutinase | 3.1.1.74 | |
| | *Rhodococcus erythropolis D4* | 1833 | | | | | 24 |
| | *Rhodococcus opacus R7* | 37919 | cutinase (Rcut) | Not specified | Abhydrolase super family | 3.1.1.74 | |
| | *Bacillus pumilus* | 1408 | Proteases, Lipases, Depolymerases | Not specified | Lipase, Depolymerase | 3.1.-.-, 3.4.-.- | 25 |

*A= EC 1.-.-.-, B= EC 1.1.1.1, C= EC 1.2.1.3, D= EC 1.10.3.2, E= EC 1.14.15.3, F= EC 3.1.-.-, G= EC 3.1.1.-, H= EC 3.1.1.3, I= EC 3.1.1.74, J= EC 3.1.1.75, K=EC 3.1.1.76, L=EC 3.1.1.101, M=EC 3.2.1.1, N=EC 3.4.-.-,O=EC 3.1.1.1*

**Supplementary Table S5.** List of plastic polymers included in the PDCOGs database (https://phylobone.com/microworld/PDCOG)

| ID | Type | Acronym | Full Name | PubChem/Other | SMILE | PSMILE | Structure | Ref |
|---|---|---|---|---|---|---|---|---|
| 1 | N | PHA | Polyhydroxyalkanoate | 405238958 | CC(O)C(=O)O | **OC(=O)C1COCN(C(=O)SSCCCCSCCOC(*)=O)CC1OCSCCCCO* | 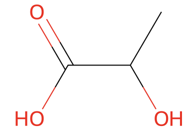 PHA | 26 |
| 2 | N | PHB | Poly(3-hydroxybutyrate) | 441 | CC(CC(=O)O)O | CC(CC(=O)O)O | 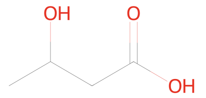 PHB | 27 |
| 3 | N | PHBV | Poly(3-hydroxybutyrate-co-3-hydroxyvalerate) | 135064954 | C(CCCC(=O)CCCC(F)(F)F)CCCBr | C(CCCC(=O)CCCC(F)(F)F)CCCBr | 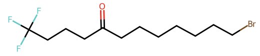 PHBV | 27 |
| 4 | N | PHBH | Poly(3-hydroxybutyrate-co-3-hydroxyhexanoate) | 464839077 | CC(O)C(=O)O ; CCCCCC(O)C(=O)O | [*]OC(=O)CH(CH3)[*].[*]OC(=O)CH(CH2CH2CH2CH3)[*] | 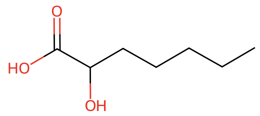 PHBH | 28 |
| 5 | N | P3HO | Poly(3-hydroxyoctanoate) | 405237853 | CC(O)C(=O)O ; CCCCCC(O)C(=O)O | [*]OC(=O)CH(CH3)[*].[*]OC(=O)CH(CH2CH2CH2CH3)[*] | 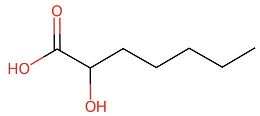 P3HO | 28 |
| 6 | N | P3HP | Poly(3-hydroxypropionate) | CN-103497922-A | CC(=O)O | [*]OC(=O)CH(CH3)[*] | 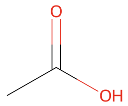 P3HP | 29 |

| ID | Type | Acronym | Full Name | PubChem/Other | SMILE | PSMILE | Structure | Ref |
|---|---|---|---|---|---|---|---|---|
| 7 | N | P4HB | Poly(4-hydroxybutyrate) | 486309711 | OCCCCC(=O)O | **OC(COC(=O)c1ccc(C(*)=O)cc1)C(OC(=O)Oc1ccccc1)C(OC(C)=O)C(=O)OCc1ccccc1* | 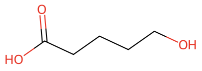<br>P4HB | 26 |
| 8 | N | PHBVH | Poly(3HB-co-3HV-co-3HHx) | Article | C(CCCC(=O)CCCC(F)(F)F)CCCBr | C(CCCC(=O)CCCC(F)(F)F)CCCBr | 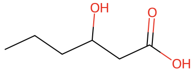<br>PHBVH | 27 |
| 9 | N | PHO | Poly(3-hydroxyoctanoate) | 405237853 | CC(O)CC(=O)O | **CC(O)COc1ccc(C2(c3ccc(OC(=O)C4(C)CC(O)C(OC(C)=O)C(C)C(OC(=O)c5ccc(O*)cc5)C4=O)cc3)C=C2)cc1* | 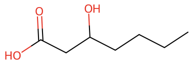<br>PHO | 26 |
| 10 | N | PHPV | Poly(3-hydroxypropionate-co-3-hydroxyvalerate) | CN-110448678-A | CC(O)C(=O)O | **CC(*)C(=O)OCC(F)(F)C(F)(F)C(F)(F)C(F)(F)C(F)(F)C(F)(F)COC(=O)C(O)COc1ccc(-c2ccc(Br)cc2S(=O)(=O)c2cccc(C(C)(C)C)c2)cc1* | 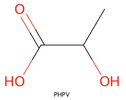<br>PHPV | 26 |
| 11 | S_He | PET | Polyethylene terephthalate | 18721140 | CC(=O)C1=CC=C(C=C1)C(=O)OCCOC | | 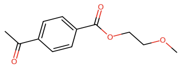<br>PET | 27 |
| 12 | S_He | PU | Polyurethane | 464858876 | CC(O)C(=O)O ; CCCCCC(O)C(=O)O | [*]OC(=O)CH(CH3)[*].[*]OC(=O)CH(CH2CH2CH2CH3)[*] | 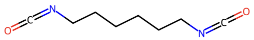<br>PU | 28 |
| 13 | S_He | PLA | Polylactic acid | 612 | CC(C(=O)O)O | CC(C(=O)O)O | 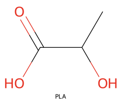<br>PLA | 27 |
| 14 | S_He | PCL | Polycaprolactone | 54078012 | CCC(COC(=O)CCCCCO)(COC(=O)CCCCCO)COC(=O)CCCCCO | CCC(COC(=O)CCCCCO)(COC(=O)CCCCCO)COC(=O)CCCCCO | | 27 |

| ID | Type | Acronym | Full Name | PubChem/Other | SMILE | PSMILE | Structure | Ref |
|---|---|---|---|---|---|---|---|---|
| 15 | S_He | PBS | Polybutylene succinate | 224143507 | CC(O)C(=O)O ; CCCCCC(O)C(=O)O | [*]OC(=O)CH(CH3)[*].[*]OC(=O)CH(CH2CH2CH2CH3)[*] | 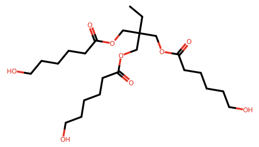 PCL | 28 |
| 16 | S_He | PBSA | Polybutylene succinate-co-adipate | 85392917 | [B-](F)(F)(F)F.CC(=O)N([CH-]C1=CC=CC=C1)C2=CC=CC=C2.C1=CC=C(C=C1)P(CCCP(C2=CC=CC=C2)C3=CC=CC=C3)C4=CC=CC=C4.[Pd+2] | [B-](F)(F)(F)F.CC(=O)N([CH-]C1=CC=CC=C1)C2=CC=CC=C2.C1=CC=C(C=C1)P(CCCP(C2=CC=CC=C2)C3=CC=CC=C3)C4=CC=CC=C4.[Pd+2] | 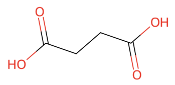 PBS | 27 |
| 17 | S_He | PES | Polyester | 481110081 | OC(=O)c1ccccc1 | **OC(=O)c1ccccc1C(=O)OCCOC(=O)/C=C(\C)C(COC(=O)c1ccccc1)C(=O)OCCCCCCOC(=O)c1ccccc1C(*)=O* | 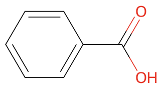 PES | 26 |
| 18 | S_He | Nylon | Polyamide | 168236 | C1CCC(=O)NCC1.C(CCCN)CCN.C(CCC(=O)O)CC(=O)O | C1CCC(=O)NCC1.C(CCCN)CCN.C(CCC(=O)O)CC(=O)O | 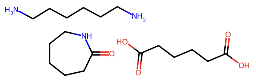 Nylon | 27 |
| 19 | S_He | PVA | Polyvinyl alcohol | 483937262 | CC(O)C(=O)O ; CCCCCC(O)C(=O)O | [*]OC(=O)CH(CH3)[*].[*]OC(=O)CH(CH2CH2CH2CH3)[*] | 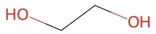 PVA | 28 |
| 20 | S_He | PBAT | Polybutylene adipate terephthalate | 85392915 | CCOP(=O)(OCC)OC1=CC=CC=C1C=CC(=NN=C(SCC2=CC=CC=C2)SP(=O)(OCC)OCC)C3=CC=CC=C3OP(=O)(OCC)OCC | CCOP(=O)(OCC)OC1=CC=CC=C1C=CC(=NN=C(SCC2=CC=CC=C2)SP(=O)(OCC)OCC)C3=CC=CC=C3OP(=O)(OCC)OCC | 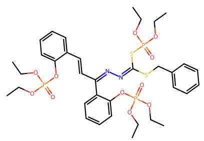 PBAT | 27 |

| ID | Type | Acronym | Full Name | PubChem/Other | SMILE | PSMILE | Structure | Ref |
|---|---|---|---|---|---|---|---|---|
| 21 | S_He | PHC | Polyhydroxycarboxylate | [US20040192881A1]() | CC(O)C(=O)O | **CC(*)C(=O)OCC(F)(F)C(F)(F)C(F)(F)C(F)(F)C(F)(F)C(F)(F)COC(=O)C(O)COc1ccc(-c2ccc(Br)cc2S(=O)(=O)c2cccc(C(C)(C)C)c2)cc1* | 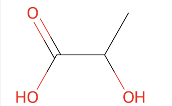 PHC | 26 |
| 22 | S_He | PC | Polycarbonate | [481110073]() | CC(O)C(=O)O ; CCCCCC(O)C(=O)O | [*]OC(=O)CH(CH3)[*].[*]OC(=O)CH(CH2CH2CH2CH3)[*] | 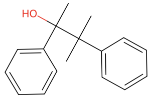 PC | 28 |
| 23 | S_He | PEG | Polyethylene glycol | [174]() | C(CO)O | C(CO)O | 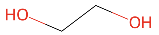 PEG | 27 |
| 24 | S_He | PEA | Polyethyl acrylate (or Polyester Amide) | [8821]() | CCOC(=O)C=C | CCOC(=O)C=C | 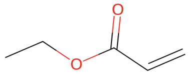 PEA | 27 |
| 25 | S_He | PEF | Polyethylene furanoate | [US-2023167232-A1]() | CC(O)C(=O)O ; CCCCCC(O)C(=O)O | [*]OC(=O)CH(CH3)[*].[*]OC(=O)CH(CH2CH2CH2CH3)[*] | 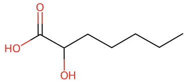 PEF | 28 |
| 26 | S_He | PMCL | Poly(ε-methylene caprolactone) | [US-9533027-B2]() | OCCCCCC(=O)O | **Oc1ccc(CC(NC(=O)COc2ccc(OC(=O)CCCCC(*)=O)cc2)C(=O)OCC)cc1OC(=O)CCCCCC* | 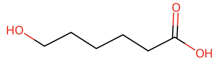 PMCL | 26 |
| 27 | S_He | Ecovio-FT | BASF blend (mainly PBAT + PLA) | [Article]() | OCC(O)C(=O) | **OC(=O)C=CC(=O)CCCCCCCCCCCCCCCCCCCC(=O)Oc1ccc(*)c(CCCCSC)c1* | 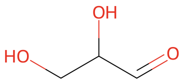 Ecovio-FT | 26 |
| 28 | S_He | P(3HB-co-3MP) | Poly(3-hydroxybutyrate-co-3-methylpentanoate) | [Article]() | CC(O)C(=O)O | **CC(C)(C)COC(=O)C(CCCCCCOc1ccc(-c2ccc(C)cc2)cc1)CC(OC(=O)CCCCC(=O)O*)c1ccc(/N=N/c2ccc(C#N)cc2)cc1* | 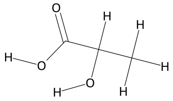 P(3HB-co-3MP) | 26 |

| ID | Type | Acronym | Full Name | PubChem/Other | SMILE | PSMILE | Structure | Ref |
|---|---|---|---|---|---|---|---|---|
| 29 | S_He | O-PVA | Oxidized polyvinyl alcohol | 405234323 | CC(=O)CO | **OC(=O)OCCCCCOc1ccc(C(=O)CCCCCCCCC(=O)CCCCCCC2C=CC2(*)O)cc1* | 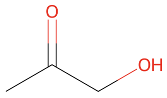 O-PVA | 26 |
| 30 | S_He | PBSet | Polybutylene succinate-co-ethylene terephthalate | Article | OC(=O)CCCCC(=O)O | **O[Si](C)(C)CCCCCCOC(=O)C(CBr)C1C(=O)OC(=O)CC1(C)C(=O)OC1COC2C(*)COC12* | 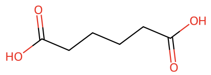 PBSet | 26 |
| 31 | S_He | Impranil | Commercial waterborne polyurethane dispersion | 472213772 | O=C=NCCCCCCN=C=O | **O[Si](C)(C)CCCCCCOC(=O)C(CBr)C1C(=O)OC(=O)CC1(C)C(=O)OC1COC2C(*)COC12* | 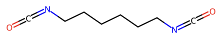 Impranil | 26 |
| 32 | S_Ho | PE | Polyethylene | 481110091 | (CH2)=(CH2) | **CCCCCCCCCCCCCCCCCNC(=O)CCCCCCCCCCCCC(=O)NCCCCCCCCCCN1CC1=CC(*)=O* | 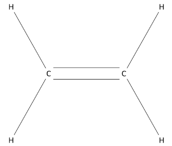 PE | 26 |
| 33 | S_Ho | PP | Polypropylene | 496176908 | CC(O)C(=O)O ; CCCCCC(O)C(=O)O | [*]OC(=O)CH(CH3)[*].[*]OC(=O)CH(CH2CH2CH2CH3)[*] | 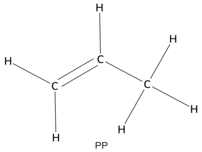 PP | 28 |
| 34 | S_Ho | PS | Polystyrene | 481110312 | CC(O)C(=O)O ; CCCCCC(O)C(=O)O | [*]OC(=O)CH(CH3)[*].[*]OC(=O)CH(CH2CH2CH2CH3)[*] | 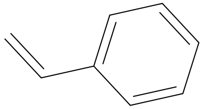 PS | 28 |
| 35 | S_Ho | PVC | Polyvinyl chloride | 481101432 | CC(O)C(=O)O ; CCCCCC(O)C(=O)O | [*]OC(=O)CH(CH3)[*].[*]OC(=O)CH(CH2CH2CH2CH3)[*] | 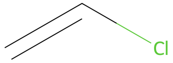 PVC | 28 |

| ID | Type | Acronym | Full Name | PubChem/Other | SMILE | PSMILE | Structure | Ref |
|---|---|---|---|---|---|---|---|---|
| 36 | S_Ho | LDPE | Low-density polyethylene | 481109052 | (CH2)=(CH2) | **CCCCCCCCCCCCCCCCNC(=O)CCCCCCCCCCCCC(=O)NCCCCCCCCCCN1CC1=CC(*)=O* | 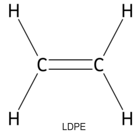 LDPE | 26 |
| 37 | S_Ho | NR | Natural Rubber (Isopren) | 6557 | CCOC(=O)C=C | CCOC(=O)C=C | 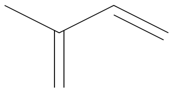 NR | 27 |
| 38 | N | P3HV | Poly(3-hydroxyvalerate) | 85305542 | CC(C1=CC=CC=C1)ON=C(C)OCC2=CC=CC=C2N(C(=O)OC)OC | CC(C1=CC=CC=C1)ON=C(C)OCC2=CC=CC=C2N(C(=O)OC)OC | 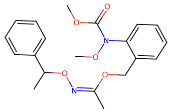 P3HV | 27 |
| 39 | S_Ho | PPL | Poly(propylene lactone) | WO2019195168A1 | CC(=C) | **CCCCCCCCCCCCCCCCCCCNC(=O)C(=O)Oc1ccc(C(=O)CCCCCCCCC(CC)(CCCCCC)C(=O)O*)cc1* | 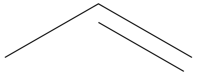 PPL | 26 |

** N: Natural; S_He: Synthetic heterochain; H_Ho: Synthetic homochain*

### SUPPLEMENTARY DATASETS

**Supplementary dataset SD1**. List of Plastic-degrading Clusters of Orthologous Groups (PDCOGs). The dataset has been created on the basis of our BLASTp result onto the NCBI COG database [30], using seed potential plastic-degrading proteins (PPDPs) from [31]. COG ID, functional categories and COG symbol were collected from https://www.ncbi.nlm.nih.gov/research/cog/ [30]. The percentage of organisms reports the ratio of total corresponding organisms (both archaea and bacteria) out of 2,296 species in the COG database. Similarly, the %archaea and the %bacteria report the amount of putative plastic-degrading bacterial and archaeal species out of total 193 and 2,103 bacterial and archaeal species respectively, in every PDCOG category. Protein_ID_PlasticDB and Enzyme_PlasticDB are annotations of the confirmed protein sequences with biodegradation capability from https://plasticdb.org/ [31].

**Supplementary dataset SD2.** Classification of polymers and plastic degrading clusters of orthologous groups (PDCOGs). Information on the PDCOGs capabilities to degrade polymer types listed both in PlasticDB [31] and others from recent literature (see Table S1). It also describes polymer properties (Natural/Synthetic and Homochain/Heterochain). The letters A-N denote putative EC classes and PDCOGs as potentially involved in the biodegradation according to recent reports (see supplementary dataset SD3). **Footnote**: 0= No, 1= Yes (putative), Letters A to N indicate enzymes with predicted potential to degradation of the plastic polymer. A= EC 1.-.-.-, B= EC 1.1.1.1, C= EC 1.2.1.3, D= EC 1.10.3.2, E= EC 1.14.15.3, F= EC 3.1.-.-, G= EC 3.1.1.-, H= EC 3.1.1.3, I= EC 3.1.1.74, J= EC 3.1.1.75, K=EC 3.1.1.76, L=EC 3.1.1.101, M=EC 3.2.1.1, N=EC 3.4.-.-

**Supplementary dataset SD3.** Matrix of organisms with the potential capacity of plastic biodegradation. Best blast hits of potential plastic-degrading proteins (PPDP) and proteins from organisms in the database of clusters of orthologous groups (COGs).

**Supplementary dataset SD4.** Metabolic characterization of plastic degrading clusters of orthologous groups (PDCOGs). This dataset includes a comprehensive characterization of the PDCOGs with information from the KEGG database [36], COG database [30], Interpro database [37], Gene Ontology database [38].

**Supplementary dataset SD5.** Environmental distribution of Plastic-degrading clusters of orthologous groups (PDCOGs) by Environments.This dataset contains information about the environments and the protein counts of all PDCOGs.

**Supplementary dataset SD6.** Observed and expected distribution of potential plastic-degrading proteins from Plastic-degrading clusters of orthologous groups (PDCOGs) database (https://phylobone.com/microworld/PDCOG). Results of the chi-squared test on the distribution of the PDCOGs by environments. The observed protein counts were taken from the supplementary dataset SD6. The results of the chi-squared test ($p$-value $< 0.05$) indicate whether the PDCOGs are present more, equal (“=”) or less than expected in a specific environment.
